# A processing pipeline for image reconstructed fNIRS analysis using both MRI templates and individual anatomy

**DOI:** 10.1101/2021.01.14.426719

**Authors:** Samuel H. Forbes, Sobanawartiny Wijeakumar, Adam T. Eggebrecht, Vincent A. Magnotta, John P. Spencer

## Abstract

**Aim:** We demonstrate a pipeline with accompanying code to allow users to clean and prepare optode location information, prepare and standardize individual anatomical images, create the light model, run the 3D image reconstruction, and analyze data in group space.

**Approach:** We synthesize a combination of new and existing software packages to create a complete pipeline, from raw data to analysis.

**Results:** This pipeline has been tested using both templates and individual anatomy, and on data from different fNIRS data collection systems. We show high temporal correlations between channel-based and image-based fNIRS data. In addition, we demonstrate the reliability of this pipeline with a sample dataset that included 74 children as part of a longitudinal study taking place in Scotland. We demonstrate good correspondence between data in channel space and image reconstructed data.

**Conclusions:** The pipeline presented here makes a unique contribution by integrating multiple tools to assemble a complete pipeline for image reconstruction in fNIRS. We highlight further issues that may be of interest to future software developers in the field.

**Significance:** Image reconstruction of fNIRS data is a useful technique for transforming channel-based fNIRS into a volumetric representation and managing spatial variance based on optode location. We present a novel integrated pipeline for image reconstruction of fNIRS data using either MRI templates or individual anatomy.

## 1 Introduction

In recent years there has been a rise in the usage of functional Near Infra-Red Spectroscopy (fNIRS) in neuroimaging studies(1–5). fNIRS is a safe, non-invasive method of neuroimaging, which measures the concentration of oxygenated and deoxygenated hemoglobin. fNIRS has become an important neuroimaging paradigm for developmental neuroscience and other fields where participant characteristics, naturalistic study paradigms, or costs make functional MRI difficult to use, such as in low-resource settings (6–8).

A key limitation of fNIRS is that optode locations can vary, making it difficult to compare across participants within a study as well as across studies. Spatial variance in brain sampling can occur due to slight differences in cap placement on the head anatomy. But even with precise cap placement, variance can occur due to subtle differences in head shape and size. This makes it challenging to make robust inferences across participants from analyses conducted in channel space given potentially erroneous but strong assumptions about the consistent placement of source-detector pairs relative to functional brain areas.

To overcome this limitation, there have been key innovations in image-reconstructed fNIRS which uses a head model as a spatial prior to transform channel-based fNIRS data into a volumetric representation. Such approaches use either Monte Carlo simulations of photon migration in the brain (9), or solve the photon diffusion equation (10), to move statistical analyses from channel space on the surface of the head into voxelspace within the brain volume. Using such methods, therefore, localizes patterns of activation from fNIRS into specific brain regions, allowing for more robust comparisons across participants and groups as data are registered into a common space. In image-reconstruction approaches, the head model used for reconstruction typically comes from an anatomical head atlas(1,11–14), although one can also use subject-specific anatomy from an individual MRI scan(15).

To date, the majority of image-reconstruction approaches have used anatomical templates, where a template atlas is used for registering the digitization and transforming into subject space. While this allows for uniformity and consistency between subjects, it overlooks the role that individual anatomical differences may play in the analyses. A subject with higher-than-average CSF in a particular area of the brain, for example, will have different tissue properties in that region which will lead to different amounts of scattering and absorption of near infra-red light. The use of individual anatomy allows such individual-level differences to inform the analyses. However, standardizing and preparing individual anatomies adds layers of complexity to the analysis -- a hurdle which is addressed in the present article.

The process of image reconstruction, particularly when using subject-specific anatomy is complex and requires multiple steps. Optode locations must be collected and checked relative to a template, with a method for correcting any digitization errors. The digitized positions must then be formatted for use with software that can create a light model. Individual MRI images must be optimized through removal of background noise, and the image must then be segmented to create the tissue model needed for photon migration simulations. The segmented head and optode locations then need to be aligned and photon migration simulations run to create the sensitivity profiles needed for image reconstruction. The fNIRS data then needs to be pre-processed in channel space and combined with the sensitivity profiles for image reconstruction. These subject-specific results then need to be analyzed at the individual level (e.g., using a general linear modelling approach) and aligned in a common group space for statistical analysis.

A current difficulty for this type of work is the lack of analysis pipelines that pull all of these steps together. No current software or published pipeline covers all aspects of this approach, making it difficult for users to have an integrated approach that includes image reconstruction within the full scope of the processing pipeline. The present article, therefore, presents a pipeline that takes the user from the initial pre-processing and cleanup steps all the way through to group analysis, with the hope that this will lead to greater transparency of results and make data sharing easier between research groups. The tools employed in this pipeline are all publicly available online.

### 1.1 Overview of the Pipeline

Figure 1 shows the steps to be followed using this pipeline. These steps are detailed below. Briefly, the first step handles preparing and cleaning the digitization data, that is, estimates of the 3D positions of the fNIRS optodes on the head. Next, we discuss how to prepare the MRI images. Here we focus on preparing individual MRI images (see left, top in Figure 1). For studies using an atlas that already contains a segmented head (see right, top in Figure 1), the user can skip to setting up and running Monte Carlo simulations to create the light model. Where users have a template MRI which has not yet been segmented, they may wish to use the segmentation steps described in this pipeline, but bypassing the cleaning steps, similar to a user in possession of individual MRIs without background noise in a good orientation. Nonetheless it is recommended that users read the steps on preparing individual anatomical files to determine whether their files meet the formatting requirements.

**Figure 1.**
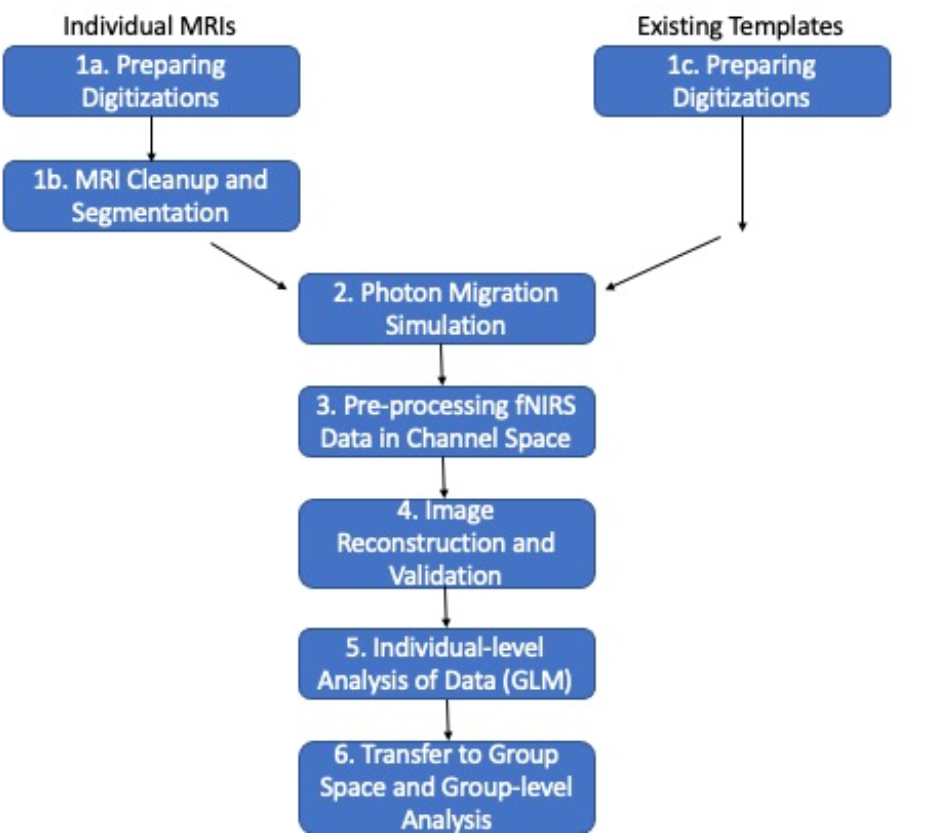
A flow diagram demonstrating the use of this pipeline. Steps on the left indicate steps to be taken with individual MRI images, steps on the right indicate steps to be taken using an existing template image.

Once the MRI data are prepared, the next step is to create the light model. There are multiple software packages that handle this step; we describe one implementation using Homer2 (https://homer-fnirs.org/) (16). Then, fNIRS data must be pre-processed. Again, there are multiple options here, and we describe a solution using Homer2. The next step is image reconstruction. Here, we use tools from the NeuroDOT package (17) (https://github.com/WUSTL-ORL/NeuroDOT_Beta). We then run an individual-level GLM on the data, again, using NeuroDOT tools. Finally, we describe how to transfer the data into a common group space for a group-level analysis. We demonstrate the final data produced by the pipeline using recent data from a study looking at the development of visual working memory in children.

## 2 Preparing Digitizations

### 2.1 Collecting digitization data

To locate the optodes on the head, we collect digitizations with a Polhemus Patriot FCC Class B digitizing device (https://polhemus.com/scanning-digitizing/digitizing-products/). The device consists of an electromagnetic receiver, placed on top of the head, and a wand used to mark the placement of the tip at each optode. The device default is set to capture the locations in inches from the receiver, but we changed this to centimeters to be compatible with Homer2 (14,16) (which we use in subsequent pipeline steps). Typically, digitizations are assumed to be in the format of one point per row, where the first set of points are reference landmarks on the head, followed by the sources, and then the detectors. Figure 2 shows a typical digitization captured with a small amount of error, pulling points out of alignment (in this case due to reaching across the head).

**Figure 2.**
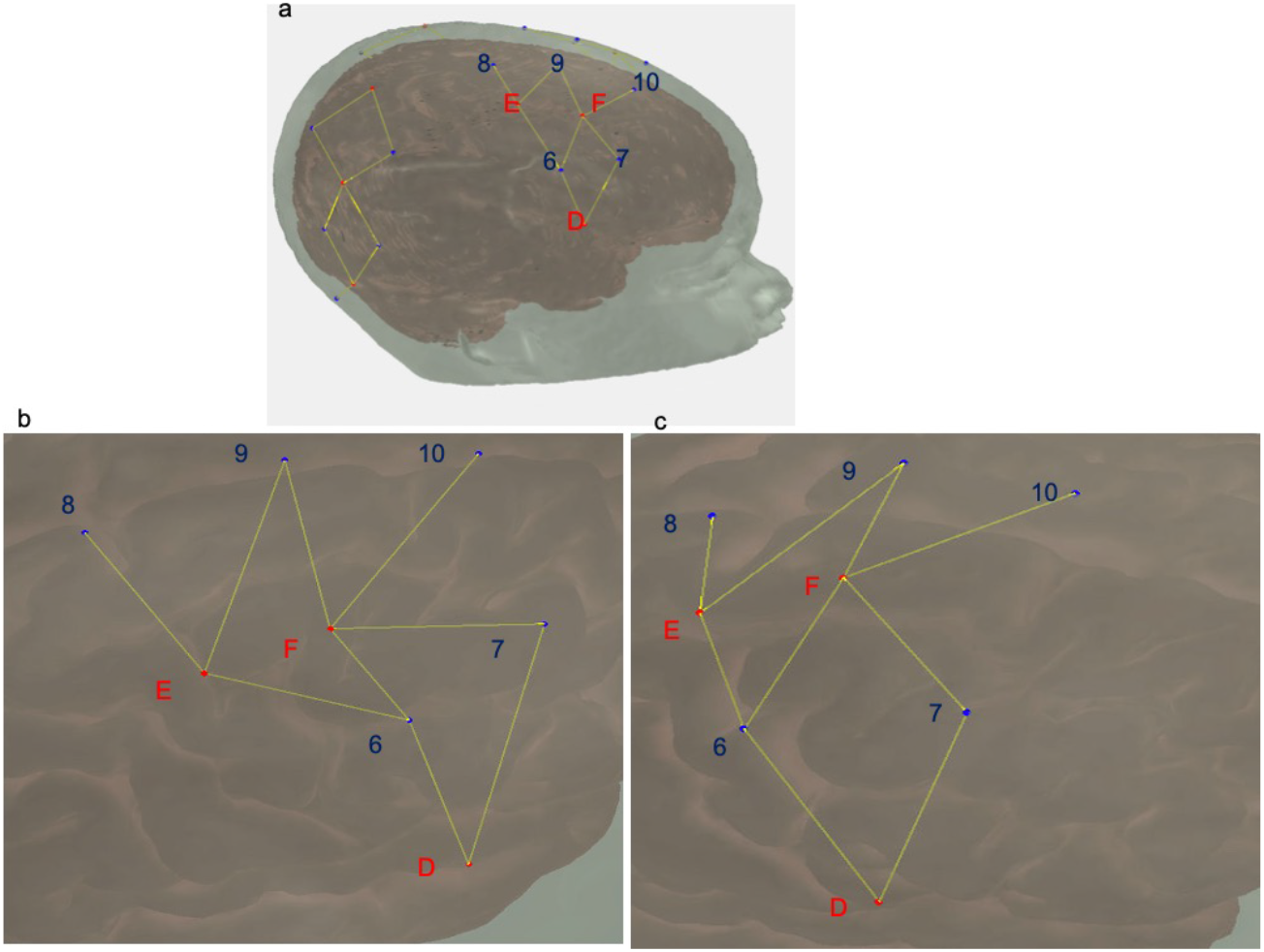
(a) The full cap layout. (b) A standard digitization with some error on the part of the experimenter, where the area shown is the highlighted region of (a). Detectors 6 and 7 (blue) have been digitized out of alignment with sources D, E, and F (red). c. The same digitization following 3-point correction. Note that the layout in (c) is the expected layout, matching the highlighted region in (a).

### 2.2 Creating digitization templates

One of the challenges of using a digitizing pen is that errors in positioning can arise. For instance, in Figure 2, several points are slightly mis-localized resulting in a distorted geometry. This can be caused by a slight slippage in the digitizing pen when the participant moves, inducing a slight mislocalisation of the pen tip. Such minor errors are difficult to identify on the fly, particularly when the head being digitized is on the body of a squirmy child. To combat errors in mis-localisation of points, we developed a package for the R language (https://www.r-project.org/), *digitizeR* (www.github.com/samhforbes/digitizeR), which takes digitized input from multiple participants for a given head size, creates a digitization template, and identifies and fixes mis-localisation errors in individual caps.

In particular, once collected, the digitizations for each participant can be read into digitizeR by cap size. For ease of visualizing caps as they change, we also recommend RStudio. The digitizeR package will read in the Polhemus output files (saved in ASCII text format) and create a list for each cap size. Next, digitizeR finds the best caps to align the other caps to for each cap size. This is done using the Kabsch algorithm (18), a method of finding the optimal rotation between two matrices to reduce the root mean squared deviation. DigitizeR uses this algorithm to align each cap to all the other caps in that size. Since every digitization runs the risk of participant movement throwing off the values for any given point (or the entire cap), each cap is rotated to match every other cap. The rationale here is that all the caps that were digitized well will all align with each other, and the caps that were digitized poorly will fail to align. So, for example, for three caps A, B, and C, if caps A and B are good caps, and C is a poor digitization, A will align with B within a given 3D criterion distance and B will align with A, whereas C will align with neither A nor B. The next step is to use this information about which caps align to create a template. In our hypothetical situation, two caps, A and B, align with each other within the criterion distance, so the template will be defined by calculating the 3D mean of each corresponding point in A and B. Using this process, a template cap is created for each cap size.

Figure 3 shows a template created from a group of caps. In this instance, the template digitization was created from a group of caps with a head circumference of 48cm. Caps were aligned using the above method where the 3D distance threshold was chosen as 10cm. This took the cap with the most matches at that cap size where no point was allowed to exceed a 3D distance of 10cm; in this instance the aligned caps had a mean distance of 3.91cm from the cap. The mean of each of the points across all of the matching caps was then calculated to create the template.

**Figure 3.**
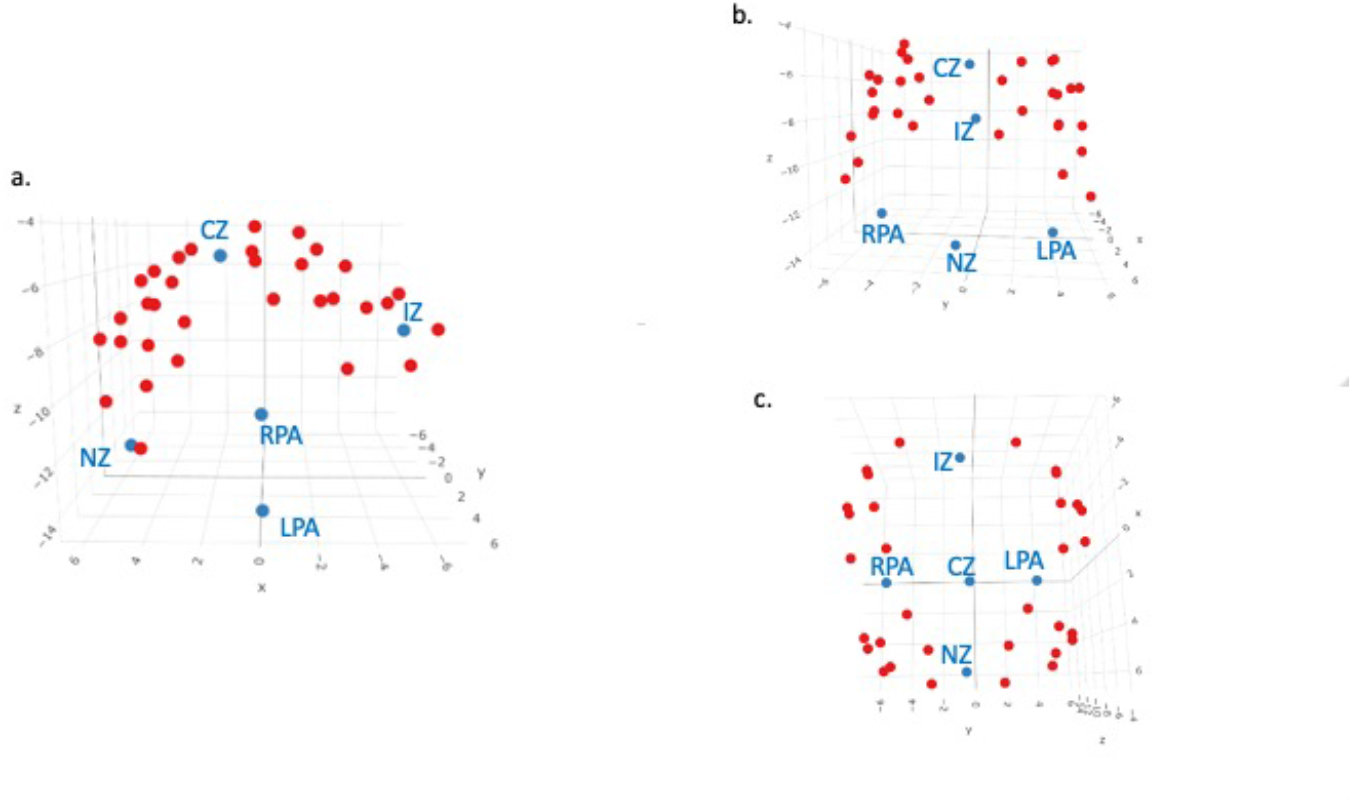
A 3D image of a template created from a group of caps, from digitizeR. Points in blue are head landmarks, points in red are digitized points from the fNIRS cap. Landmarks have been labelled. (a) shows the cap from the left-hand side, (b) from the front, and (c) from the top.

Once created, templates can be saved in digitizeR and used for all participants in that cap size if desired. This would yield an average acceptable digitization that is unique to each cap size, but captures the data collected across the sample. Alternatively, as we describe in the next section, digitizeR allows using the template to clean and save modified versions of the individual digitizations, allowing the user to preserve as much of the individual-level variation in optode location and cap placement as possible.

### 2.3 Correcting individual caps

Once templates have been constructed, each participant cap can then be aligned to the template for that cap size, again using the Kabsch algorithm. There are two sources of variation in the digitized locations in 3D space: true variation in cap placement and optode location due to factors such as wrinkles in the cap or the cap being off-center; and errors such as digitization of the incorrect point or movement causing the point to be unrealistically far away from the head. The goal of the correction at the individual cap level is to permit as much of the former variance as possible, which is due to the true placement of the optode on the head, while eliminating the large sources of the latter type of variance.

The digitizeR package allows three different correction methods: a three-point method, an iterative method, and a head-wise method. The three-point method allows the user to specify three threshold 3D distances, at which the points will be sequentially tested against the template for that cap size, and those outside those distances will be removed, and then the cap re-aligned without those points. After the final distance check, the missing points are replaced with points from the relevant template. The difficulty with the three-point method is selecting the three distances, and this can be dependent on the data. The data that were cleaned using the template in Figure 3 were cleaned using threshold distances of 12, 10 and 7cm, with the end goal of making sure no more than 1/3 of the original data points had been replaced using this method.

The iterative approach takes the same method as the three-point approach, but rather than the user-specified distances, it starts with a 3d distance of 20cm and replaces points outside of that distance, and then reduces the difference by 1cm, repeating the procedure of aligning to the template and removing the points outside of the 3D distance each time. The headwise method follows a similar logic, but rather than individual points being replaced, the mean 3D distance from all the points is calculated against the template.

In this report, we used the three-point method, although the best approach may be dependent on the types of errors being seen in the data. In the case of our dataset, we wanted to minimize the number of points we were replacing while ensuring our caps were as correct as possible. While some trial and error is necessary to determine the best distances for the caps at hand, the three-point method is a good option for this purpose. The points that were removed as part of this procedure are then filled in with the equivalent point from the template once the cap is aligned to make a complete cap that is aligned in the same space as the template cap.

### 2.4 Preparing for use with Homer2

The final versions of the digitized caps after alignment to the template can be saved and exported from R using the built-in functions from the digitizeR package. The save_caps option automatically saves each of the caps in its own folder as a digpts.txt file, saved in the format that is necessary for Homer2, which includes reference, source, or detector labels for each point. In addition, the caps are converted by default to distances in millimeters (from centimeters), making them automatically ready to use for import into Homer2. Full documentation for any of the digitizeR functions can be found at www.github.com/samhforbes/digitizeR.

## 3 MRI Cleanup and Segmentation

One of the challenges with fNIRS image reconstruction currently is that there are few pipelines that can handle both atlas-based and individual-MRI-based registration. The latter case is particularly challenging as individual MRIs require both cleanup and segmentation before this information can be used in the registration process. These steps can be quite challenging, particularly in developmental research as infant and child MRI data can have unique characteristics. Thus, in this section, we describe steps in our pipeline to help researchers standardize MRI cleanup and segmentation.

The segmentation process is based on having an anatomical T1-weighted image from each participant. The T1-weighted MRI images can be treated differently in this pipeline based on the age of the participants and image quality. The segmentation process uses tools from AFNI (Analysis of Functional NeuroImages; https://afni.nimh.nih.gov) to generate a label map that can readily be imported into AtlasViewerGUI, a visual graphical user interface contained within Homer2 (https://homer-fnirs.org/) (14,19), providing brain and skull surfaces used in the analysis of the fNIRS data. The steps of the segmentation process are outlined here along with the options that can be employed to handle scans from infants as well as low SNR images.

The starting point for the processing pipeline is an anatomical T1-weighted image that approximately has the nose aligned with the y-axis of the scanner (i.e., the nose should be pointing forward (anterior) in an axial image). If the orientation of the nose is more than 15 degrees from this orientation -- for example in a sleeping infant -- we have found it helpful to rotate the image such that the nose is roughly aligned with the y-axis for initiating the pipeline. This can be done using 3drotate from AFNI or similar functionality from other image analysis packages. Next, the image is resampled (3dresample) into a standard Right-Axial-Superior orientation. This ensures that the image will be oriented properly once imported into AtlasViewer. The next step of the segmentation pipeline is bias field correction (3dUnifize); this is optional and can be used if large variations in the signal exist across the image. We then create a brain mask to define brain tissue in the image. This step is sensitive to the age of the subject being analyzed since infants have a significantly smaller intracranial volume as compared to adults. Thus, this step has two options to generate the brain mask: 3dSkullstrip and ROBEX (20). The 3dSkullstrip option performs well in most cases and the user is provided with the ability to set the initial radius and expansion rate for the sphere inflation used in the underlying algorithm. In infant scans, we have found improved performance with the ROBEX-based brain mask. For details on using these functions, please see the instruction guide available at https://github.com/developmentaldynamicslab/MRI-NIRS_Pipeline.

The next step of the pipeline is to put the T1 weighted scan into AC-PC alignment. To do this, the brain mask in the prior step is used to extract the brain from the T1-weighted image, which is subsequently aligned with a skull-stripped Talairach Atlas image using auto_tlrc command from AFNI. The rigid body transform from the resulting transform is saved and used to reorient the subject T1-weighted scan and brain mask into AC-PC alignment. Next, we pad the image with zeros. This ensures that the surface generated will be ‘closed’ when imported into AtlasViewer. We then generate a skull mask from the image by identifying an optimal threshold (3dClipLevel) for background removal. The resulting mask generated from thresholding at this optimal level is then filled to define all voxels within the skull. The next step of the pipeline is optional and median filters the image. This may be necessary for low SNR images. The final step of the pipeline uses the 3dSeg command from AFNI to define the tissue into gray matter, white matter, and CSF. The resulting segmentation is then combined with the skull segmentation to generate a label map that contains 4 labels: gray matter, white matter, CSF, and skull.

Difficulties in this processing pipeline can arise when the original T1-weighted MRI image has a large amount of background noise, such as having captured additional objects (such as the participant’s arm) in the scan. For these more extreme cases, we have found it helpful to align the scan to a template, and then create a filled mask of the template, which is used to remove the background noise from the original scan. This leaves the image with only the head and no extraneous noise. Further information on this is provided in the pipeline instructions in the repository.

Once these steps are complete, the label map can then be read into Homer2 to create the anatomical files needed for the spatial prior (i.e., the light model). This image is read in using the import MRI functionality from *AtlasViewerGUI*, and then anatomical reference points are selected by the user. This allows for the construction of the anatomy which can then be combined with the cleaned digitizations from the prior step.

## 4 Photon Migration Simulation

The next step is to create the light model that will be used for image reconstruction. As this uses previously described tools (14), we describe these steps briefly here. We used the *AtlasViewerGUI* package within Homer2 to project the corrected digitized points for each participant onto the segmented atlas created by cleaning up and segmenting the participant’s MRI scan. To ensure that the photon migration simulations were age-specific, absorption and scattering coefficients for the scalp, CSF, gray and white matter for both wavelengths of light were provided such that each node of the mesh was assigned optical properties determined by tissue type based on values from the literature (17,21–24). Photon migration Monte Carlo simulations with 100 million photons were run to generate sensitivity profiles for each channel and wavelength. In the data presented here, this process was carried out using the High Performance Computing cluster supported by the Research and Specialist Computing Support service at the University of East Anglia, which enabled the running of simulations for all channels and participants in parallel by utilizing multiple cores. As this is computationally expensive, we recommend use of a server or other high performance computing device for this step. Next, the head volume and the sensitivity profiles were converted into NIFTI format for use in image reconstruction. Sample photon migration simulations are shown on a segmented head from the MRI scan of a 6-month old in Figure 4a. The head volume and an exemplar channel for the same participant is shown in Figure 4b.

**Figure 4.**
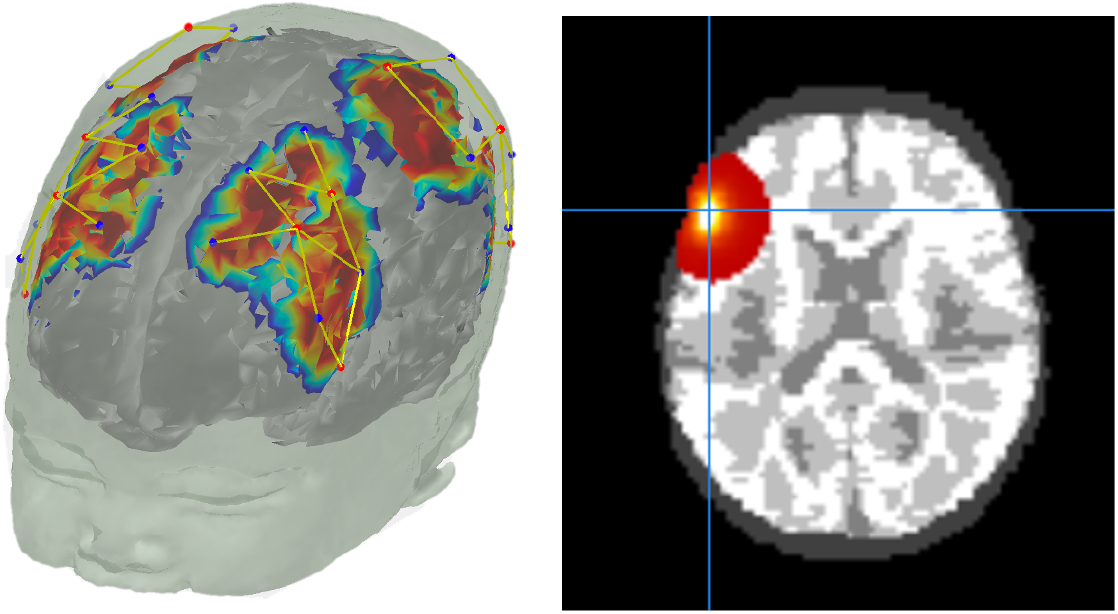
(a) Photon migration simulations on a segmented head of an MRI scan from a 6-month old infant. (b) Sensitivity profile of an exemplar channel from the same participant in NIFTI format (visualized using *Mango*)

## 5 Pre-processing fNIRS data in channel-space

For the purposes of demonstrating the quality of image reconstruction below, we used data from a TechEn CW7 system (12 sources and 24 detectors) with wavelengths of 690 nm and 830 nm. Fiber optic cables were used to carry data from the machine to a customized probe geometry of 40 channels covering bilateral frontal and parietal cortices (see Figure 4a). fNIRS data was preprocessed using the *EasyNIRS* package in Homer2. Raw data was pruned to include channels with signals between 80 dB and 130 dB using the *enPruneChannels* function. Next, these data were converted to optical density units using the *hmrIntensity2OD* function. Motion artifacts were corrected using the *hmrMotionCorrectPCArecurse* function which used targeted principal components analysis (25) [tMotion = 1.0, tMask = 1.0, StdevThresh = 50 and AmpThresh = 0.5]. The corrected data was examined for uncorrected motion artifacts using *hmrMotionArtifactByChannel* [tMotion = 1.0, tMask = 1.0, StdevThresh = 50 and AmpThresh = 0.5]. If these uncorrected artifacts fell within −1 to 18 seconds of a stimulus trigger, that trigger was removed from further data processing using the *enStimRejection* function. Finally, these data were bandpass filtered using the *hmrBandpassFilt* function with high-pass and low-pass cutoff frequencies of 0.016 and 0.5 Hz, respectively. Relevant information from these processed .nirs files were then used in subsequent steps.

## 6 Image Reconstruction and Validation

The next step in the processing pipeline is image reconstruction. This step integrates the 3D light model created using *AtlasViewer* with the channel-based fNIRS data processed using Homer2. The image reconstruction is conducted in Matlab using NeuroDOT tools (17) (https://github.com/WUSTL-ORL/NeuroDOT_Beta). In the sections below, we provide an overview of the image reconstruction approach including details for streamlining processing steps between AtlasViewer, Homer2, and NeuroDOT. We then show sample image-reconstructed data plotted relative to the channel-based fNIRS data.

### 6.1 Overview of Image Reconstruction in NeuroDOT

First, the sensitivity volumes for each source-detector pair that were calculated in *AtlasViewer* were converted into NeuroDOT format. In NeuroDOT, the sensitivity volumes are arranged in a single two-dimensional matrix *A* with measurements in the first dimension and voxels in the second dimension. A structure variable *info* contains sub-structures that house meta-data describing the sensitivity volumes (*info.tissue*), the optode locations (*info.optodes*), the measurement list (*info.pairs*), the stimulus paradigm (*info.paradigm*), and other parameters from data acquisition (*info.system*). The header that contains the spatial meta-data from the NIFTI *.nii files from AtlasViewer is contained within *info.tissue.infoT1*. The order of the measurements in *A* corresponds to that described in the measurement list located in *info.pairs*. The voxels are ordered as if the entire sensitivity volume is unwrapped into a single vector. To maximize computational efficiency, only voxels with a sensitivity above a given threshold for any of the measurements are included. The indexing of these voxels relative to the entire volume is described in the variable *info.tissue.dim.Good_Vox*. This step typically saves around a factor of 5-10x in disk space and lowers computation time due to lower demand on RAM. For this study, the threshold was set to 0.01 of the maximum sensitivity within the combined volumes of all measurement sensitivity profiles.

Second, the preprocessed optical density data from Homer2 in the *.nirs file was converted directly into NeuroDOT format. The NeuroDOT measurement list represented in the table *info.pairs* is populated from the *SD* variables in Homer2. The table *info.pairs* contains the source, detector, wavelength, source-detector separation in 2- and 3-dimensions, and other information for each measurement. The measurements to be used for reconstruction are listed in the *info.MEAS.GI* variable. The stimulus paradigm timing information is populated into *info.paradigm* for use in the post-reconstruction analyses (described below).

To minimize contributions from extra-cerebral tissue in the variance of the measurement data, we performed global signal regression separately for each wavelength. The mean signal across all low-noise measurement channels was regressed from all channels in a manner akin to superficial signal regression as previously described using the NeuroDOT function *regcorr* (26–28). Though short measurement channels were not utilized in these data, the mean across all measurement channels provides an estimate of systemic physiology present in all of the data. Removing this global mean increases the contrast to noise and spatial specificity of the variance remaining in the measurement data. The conversion of measurement-wise data to voxel-wise data through reconstruction significantly increases the overall size of the optical data due to the presence of many times more voxels than measurements. To help alleviate the large computational demands, the data was temporally down-sampled to 25 Hz before reconstruction using the NeuroDOT function *resample*_*tts*.

Once the optical, sensitivity, and meta-data have been converted into NeuroDOT format, the sensitivity matrix can be inverted separately for each wavelength using the Moore-Penrose generalized inverse with a Tikhonov regularization parameter of λ_1_=0.01 and spatially variant regularization parameter of λ_2_=0.1 using the NeuroDOT function *Tikhonov_invert_Amat* (29). The Tikhonov regularization parameter tunes the balance between spatial smoothness and high-spatial frequency noise in the image. Spatially variant regularization helps control for the tendency for DOT methods to reconstruct too superficially and is set to optimize spatial correspondence with fMRI based on other studies (30). The optical data are reconstructed into the voxelated space using the NeuroDOT function *reconstruct_img*.

Relative changes in oxygenated (HbO_2_), deoxygenated (HbR), and total hemoglobin (HbT) concentrations are obtained from the reconstructed absorption coefficients by spectral decomposition of the extinction coefficients of HbO_2_ and HbR at the two wavelengths (31,32). Conversion from relative changes in absorption coefficients to HbO_2_ and HbR is performed with the NeuroDOT function *spectroscopy_img*.

### 6.2 Sample Image Reconstructed Data

Figure 5 below shows samples of image reconstructed fNIRS data from one participant plotted against the processed channel-based fNIRS data. To create these images, we first placed the centroid of a 2cm diameter sphere at the maximum value of each channel’s sensitivity volume from the photon migration simulations for each chromophore. Note that we constrained the sensitivity volumes to the cortex to ensure that the maximum value was located in the cortex and not in the skull or surface tissues. Next, we extracted the mean reconstructed fNIRS time series from each sphere, averaging over voxels within the sphere. The red curve in each panel shows the mean image-reconstructed time series, while the black line shows the channel-based data from the associated channel. We show data from 6 channels in total (see brain images in Figures 5E, F for mapping of channels to spheres). We selected these channels because they are located in parietal and ventral occipital regions of interest for our sample data set. As can be seen in the figure, the image-reconstructed data match the channel-based data well for both HbO_2_ (Figure 5A,B) and HbR (Figure 5C, D). The signals are within the same range, with the image-reconstructed data typically showing a bit more signal intensity.

**Figure 5.**
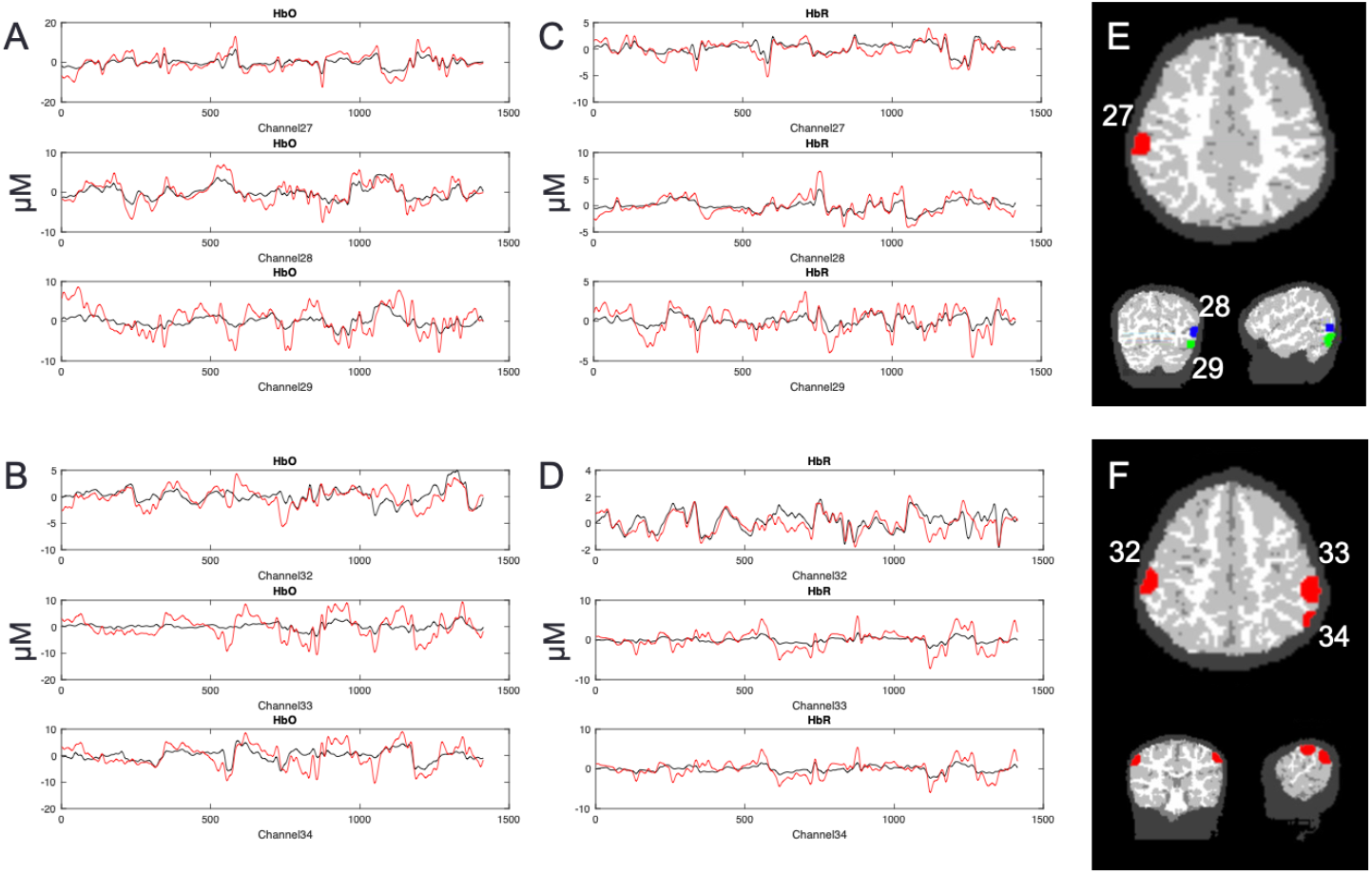
Image reconstructed fNIRS data (red lines) plotted against channel-based fNIRS data (black lines). Left panels show HbO time series (in μM) for channels 27-29 (A) and channels 32-34 (B); middle panels show HbR time series for channels 27-29 (C) and channels 32-34 (D). Panel E shows the locations of the spheres used to extract data for channels 27-29. Note that different colors are used to show the channels across views in E as the axial slice does not match the coronal and sagittal views. Panel F shows the locations of the spheres associated with channels 32-34 (axial, coronal, and sagittal views are all aligned).

### 6.3 Validating the Image Reconstruction Approach

To validate the image reconstruction approach, we correlated the average image-reconstructed time series (e.g., red lines in Figure 5) with the channel-based time series (e.g., black lines in Figure 5) for all data runs for four sample participants. In total, we conducted 504 correlations: 7 runs (over 4 participants) x 36 channels x 2 chromophores (HbO_2_/HbR). Data from 18 channels were discarded because the channels were pruned in the initial fNIRS processing. The correlation results are shown in Figure 6. Most of the correlations were quite high and above our minimum acceptable threshold of 0.25. In particular, 473 of 496 correlations were greater than 0.25; the mean *r* value for this subset was 0.8. Thus, only 2.6% of the sample was below the criterion. In these cases, the channel-based fNIRS data was of low quality with poor SNR.

**Figure 6.**
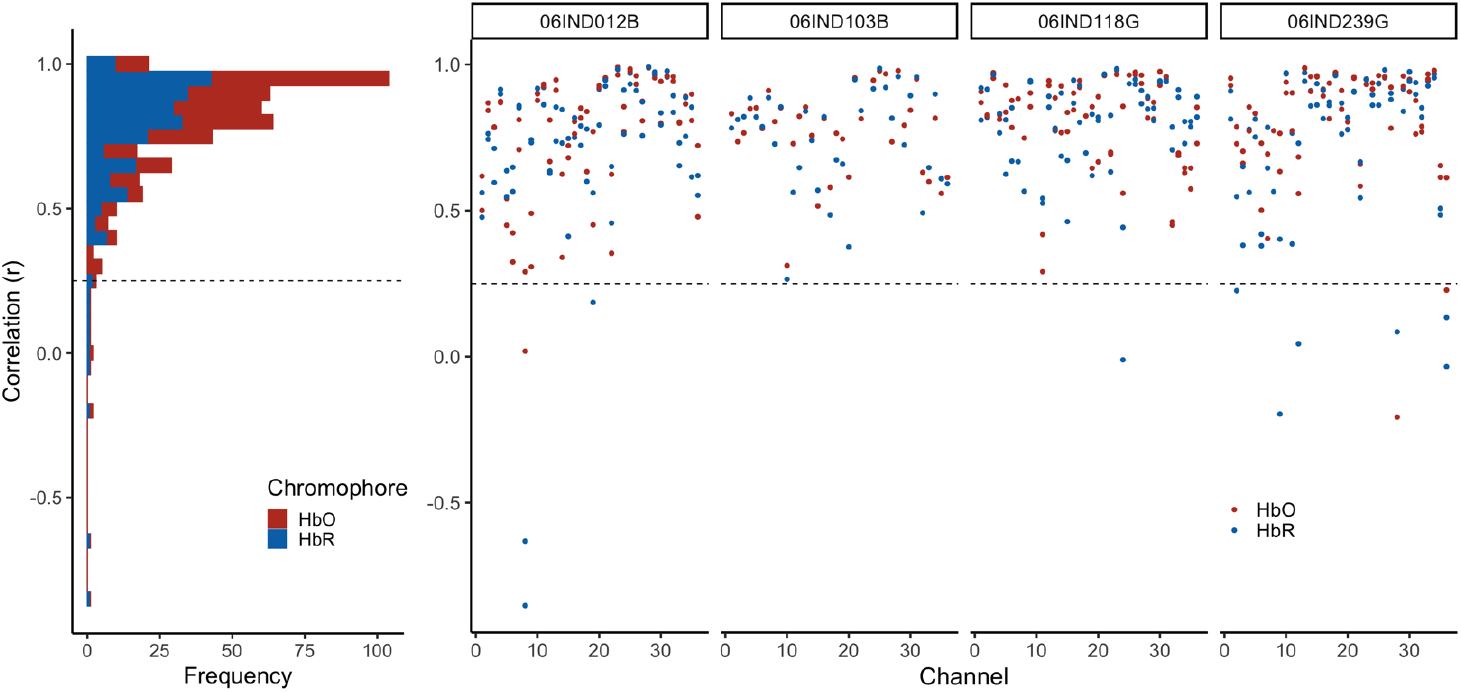
Validating the fNIRS image reconstruction data. Each dot shows the correlation between the image-reconstructed fNIRS data averaged over a 2cm sphere in cortex (see examples in Figure 5E,F) and the associated channel-based data. Red dots show correlations for HbO; blue dots show correlations for HbR. Data are shown for four participants (see columns). Three participants (06IND012B, 06IND118G, 06IND239G) had multiple runs, so there are four correlation values per channel (HbO/HbR for run1 and run2). Dashed line shows our target criterion value of 0.25. The vast majority of comparisons were above the criterion value.

## 7 Individual-level Analysis of Data (General Linear Model)

Once the participant fNIRS data have been reconstructed, a typical approach is to model these time series data using general linear modelling. For our test dataset, brain responses were analyzed using a general linear model (GLM) run with the NeuroDOT function *GLM_181206*. We used a hemodynamic response function (HRF) derived from DOT data as it has been shown to be a better fit for optical data than those derived from fMRI (33). The same HRF was used for HbO_2_ and HbR responses. Each event was modelled with a 10 second extent and variable intertrial intervals with an exponentially distributed set of durations. We focused our analyses over the window from one second before each stimulus onset to 20 seconds post onset. To control for variability in the number of events per run and per participant, we computed a weighted average of the betas over runs, weighted by the number of events per condition in each run.

## 8 Transfer to Group Space and Group-level Analysis

This step can employ any common or group space that may be applicable for the study. Often it is convenient to use the MNI coordinate system to report functional activation since it is the standard space often used for fMRI and PET activation studies. The registration from subject space to the atlas space is driven by the AC-PC aligned image and brain mask generated in Section 3 and corresponding T1-weighted image and brain mask of the template image for the group space. We used ANTS software to implement a hierarchical image registration procedure that includes a high dimensional diffeomorphic non-linear symmetric image normalization (34) as the last step in the registration. The prior steps of the image registration hierarchy include a rigid body registration followed by an affine registration. The resulting transform was then applied to the HbO_2_ and HbR beta maps from the individual-level GLM results.

## 9 Demonstration of the pipeline on an example dataset from NIRSSport (NIRX)

To demonstrate the type of data the processing pipeline yields, we report sample results here from a second test dataset taken using a NIRSSport (NIRx) system. This has the added utility of showing that our pipeline generalizes well across NIRS systems (as the prior examples in this report used a TechEn system). The data presented in this case study are part of a longitudinal study investigating the effects of schooling on neurocognitive function (35).

### Participants

Data from seventy-four 4.5-year-old children (34 females, *M* age = 53.5 months, *SD* =1.3) who participated in a longitudinal study in Scotland were used to validate the analyses pipeline. The study was conducted in the participants’ homes. Parents gave written informed consent and children gave verbal assent prior to the testing session. The research was approved by the General University Ethics Panel (GUEP 375) at the University of Stirling.

### Experimental Task

Visual working memory (VWM) performance was measured as children took part in a color change detection task (36). The experimenter explained the task to the children using 3×3 inch flash cards. Once the experimenter was satisfied that the children understood the rules, they moved to the experiment. The experimental task was run in E-prime V.3 software on an HP laptop with a 14-inch screen. In each trial, a fixation appeared on either side of the screen to orient the child to the screen. Then, a memory array of colored squares was presented for 2 seconds, followed by a blank screen for 1 second (delay period). Then, a test array of colored squares was presented and stayed on the screen until a response was made. After the presentation of the test array, the child was asked whether the colored squares were ‘same’ or different’ between the memory and test arrays. The experimenter registered the child’s response using the keyboard. Each trial was followed by an inter-trial interval of 1 second (50% of the trials), 3 seconds (25% of the trials), or 5 seconds (25% of the trials). VWM load was manipulated from 1 to 3 items (load 1, load 2 and load 3). There were 8 trials for the same condition and 8 trials for the different condition for each load. Thus, there were 12 possible conditions for trials in this task, 3 loads (1, 2 and 3) x 2 trial types (same and different) x accuracy (correct and incorrect).

### fNIRS Data Acquisition

A NIRSport system (NIRX) with 8 sources and 8 detectors was used to collect neuroimaging data from children as they engaged in the VWM task. Data were collected at 7.81 Hz and the wavelengths of the sources was 760 nm and 850 nm. Probe geometry with fourteen channels overlaying the bilateral frontal and parietal areas (see Figure 7a) was constructed based on regions of interest from previous fMRI VWM literature (37). The head circumference of the child was measured and a suitable cap was chosen. Four cap sizes were used across all children and source-detector distances were scaled according to cap size (50 cm: 2.5 cm, 52 cm: 2.6 cm, 54 cm: 2.7 cm and 56 cm: 2.8 cm). The distances from the nasion to inion and between the peri-auricular points were measured to ensure that the cap was centrally placed on the child’s head.

**Figure 7.**
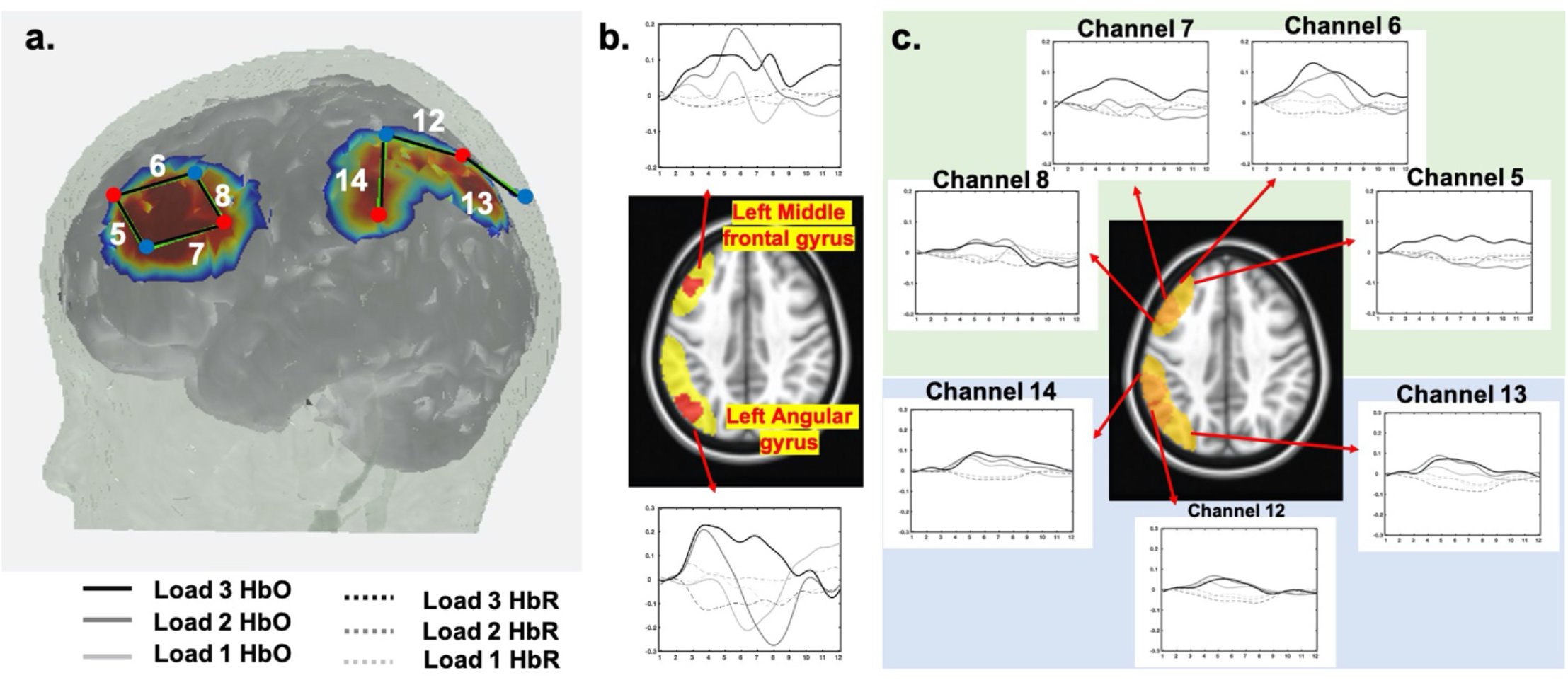
a. Probe geometry for the task. b. Significant clusters from the image-reconstructed fNIRS data in the middle frontal gyrus and left angular gyrus (see red clusters in brain image). Time series plots show weighted average HbO_2_ (solid lines) and HbR (dashed lines) for load 1 (light grey), load 2 (medium grey), and load 3 (black) conditions. c. Block-averaged activation from the channel-based fNIRS data showing channels overlapping with the red clusters in (b) (see highlighted yellow region in brain image). Plotting conventions as in (b).

### fNIRS Data processing

The probe geometry was digitized to obtain the coordinates of sources and detectors for each of the four capsizes. A 4.5-year-old MRI atlas was obtained from the Neurodevelopmental MRI database (38–42) and segmented into four tissue types (scalp, CSF, grey matter and white matter). Monte Carlo simulations were run with 1 million photons to generate sensitivity profiles for channels for each capsize (step 2, see Figure 1). The head volume and sensitivity profiles were converted into NIFTI format. Next, channel-based fNIRS raw data was pre-processed, corrected for motion artifacts, filtered, and converted to optical density units. Data were also corrected for superficial scalp artifacts using global signal regression (step 3). These channel-based fNIRS optical density data was integrated with the volumetric sensitivity profiles using image reconstruction as described previously (step 4).

A general linear model with 6 regressors (for 12 conditions) was run on each voxel by convolving a modified gamma function from SPM (delay of response = 4; delay of undershoot = 15, dispersion of response = 1; dispersion of undershoot = 1; ratio of response to undershoot = 6; onset =0; length of kernel =16) (43) with a boxcar window of duration 4.5 seconds (step 5). Note that this gamma function was chosen after visually comparing the duration of response onset, time to peak and response offset of the generated HRFs and conventional channel-based block averages. Further, this boxcar window was chosen to account for a sample array of 2 seconds, delay of 1 second and the rest of the time dedicated towards the test array period. Beta coefficient maps were obtained for all 12 conditions in each voxel for each chromophore and participant. Next, these individual-level beta maps were registered to the MNI space (step 6). For Group analyses, the beta maps were entered into a linear mixed effects model (using *3dLME* function in AFNI) with within-subjects factors of load, trial type, and chromophore (HbO and HbR). Note that only correct trials were used for our analyses.

For the current case-study, we focused on the interaction between load and chromophore to examine whether HbO activation in significant clusters show the canonical trend of increasing activation with increasing VWM load. Only those voxels that contained data from 60% of the subjects were included for further analyses. To control the family-wise error in our data, we used the 3dFWHMx function in AFNI to estimate the empirical ACF in our fNIRS data and fit a mixed ACF model to this function. We then used the mixed-ACF parameters (0.9939 8.3354 5.2485) in 3dClustSim with a voxelwise *p* = 0.01, alpha = 0.05, and 10,000 iterations. We selected two-sided thresholding with the NN1 option (first-nearest neighbour clustering). The cluster size criterion was 169 voxels.

### Results

Two clusters showed a significant interaction between load and chromophore: left middle frontal gyrus and left angular gyrus. Figure 7b shows the two clusters in red. Block averaged change in HbO/HbR activation extracted from these clusters are also shown. In the left angular gyrus, HbO activation increased with increasing load. In the left middle frontal gyrus, activation was highest for load 2. Activation in both regions show the canonical difference between HbO and HbR activation, with the largest difference visible at the higher loads. Masked sensitivity profiles from channels 5, 6, 7 and 8 in the left frontal cortex and 12, 13 and 14 in the left parietal cortex are shown in yellow. Block-averaged activation from these channels are shown in Figure 7c. We see similar trends in activation with increasing load between the block averages from the significant clusters from the image reconstruction approach and block averages from the conventional channel-based approach. Image reconstruction provides more precise localization of the source of activation in the brain. The change in activation with increasing load is more pronounced when localized to the left angular gyrus compared to the changes in activation in the three channels. It is important to note that the goal of employing image reconstruction techniques is not to empirically compare against and arrive at the same result observed in channel space. Instead, it affords the opportunity to obtain spatial localization of activation from multiple channels to specific clusters and compare these activation maps across participant groups and ages while accounting for key sources of spatial variance in the data.

## 10 Discussion/Conclusion

Image reconstruction of fNIRS data is a critical tool in standardizing the spatial location of fNIRS data from both individual and multiple channels into voxel space. Doing so allows for analysis of fNIRS data based on anatomical region rather than channel, accounting for several known sources of spatial variance in the data including variance in exact optode location on any given participant. This approach also enables users to tap powerful fMRI analysis tools in the process. Despite the considerable benefits of this approach, there are complexities in the analysis that make image reconstruction difficult. These include a lack of standard approaches for cleaning digitized optode locations and creating templates, standardizing individual anatomies, creating a light model based on either template or individual data, running the general linear model, moving the data into a common space, and analyzing the data. In the present article, we described a robust pipeline with accompanying code to push through these complexities along with initial validation data acquired from different fNIRS systems.

In the present report, we demonstrated that with fNIRS data and both individual anatomical MRI images and templates, fNIRS data can be reconstructed in a manner that is faithful to the original data, and analyzable in voxel space. The accompanying publicly available code can be adapted by users for their own data requirements. The code also contains additional functionality not discussed in the current paper, such as functions to cross-check GLM results against the timecourse data from a given region. We also demonstrated with datasets from child fNIRS studies that the analyses and reconstruction methods are reliable and valid.

The pipeline presented here contains several important contributions to the field. In the *digitizeR* package for the R language, we present a novel method of creating digitization templates and preparing them for use with other software. In addition, we provide code, suggestions and guidelines for the cleaning and segmenting of individual MRIs to turn individual anatomical data into a brain model suitable for use in the creation of a light model. We then used a combination of existing software to demonstrate how to create a light model, perform a general linear model, and move data into group space for analysis. While some individual aspects of this pipeline, such as the creation of a light model, feature in a number of software packages, these have not previously been combined in a way that allows users to go from raw data to final group analyses as reported here. The pipeline reported here, therefore, provides a roadmap of the functionality that users require to take their data all the way through analysis. As such, we hope this paper will be of use to software developers and others interested in creating standardization of analysis approaches within a rapidly developing field.

## Disclosures

The authors have no competing interests to disclose.

## Acknowledgements

This work supported by Grant No. OPP1164153 from the Bill & Melinda Gates Foundation and Grant No. R01HD083287 from the National Institutes of Health. Both grants were awarded to J. P. Spencer. The authors are grateful for the contributions of Aaron Buss and Jessica Defenderfer for their comments during testing of the pipeline.

## Notes

### Competing Interest Statement

The authors have declared no competing interest.

